# FBH1 restrains replication progression independently of promoting double strand breaks at hydroxyurea-stalled replication forks

**DOI:** 10.1101/2025.07.08.663736

**Authors:** Joshua L. Turner, Georgia Moore, Samantha L. Reed, Tyler J. McCraw, Zane R. Tolbert, Jennifer M. Mason

## Abstract

In response to damage, stalled replication forks undergo fork reversal converting replication forks into a 4-way junction. FBH1 is a DNA helicase that promotes replication fork reversal. After prolonged replication stress, FBH1 promotes double strand break accumulation and cell death. One of the hallmarks of loss of fork reversal is unrestrained replication in the presence of DNA damage, but how FBH1 restrains replication at stalled forks is not known. Here, we show that FBH1 limits replication fork progression by PRIMPOL and REV1 in response to hydroxyurea. However, neither PRIMPOL nor REV1 is preventing replication fork collapse in FBH1-deficient cells. We demonstrate that increased resistance of FBH1-deficient cells to hydroxyurea treatment is not due to mis-regulation of PRIMPOL. Surprisingly, REV1 depletion sensitized FBH1-deficient cells to hydroxyurea without restoration of double strand break induction. Together, our results indicate FBH1 restrains replication and promotes replication fork collapse by independent mechanisms.

## Introduction

Replication forks encounter impediments that slow or stop replication fork progression threatening genome integrity. Prolonged replication fork stalling results in double strand break formation (i.e., replication fork collapse). To protect the integrity of DNA during replication, cells have evolved complex mechanisms involving DNA damage signaling, DNA repair pathways, and DNA damage tolerance (DTT) pathways collectively referred to as the replication stress response. These mechanisms resolve any underlying damage and allow for the completion of replication (1).

Replication fork reversal occurs when a 3-way junction is converted into a 4-way junction by pairing of nascent strands (2). Two independent fork reversal pathways exist in mammalian cells, one pathway is mediated by HLTF, SMARCAL1, and ZRANB3 and a second pathway is mediated by FBH1 (3–7). Both fork reversal pathways require RAD51 and RAD54, however, only the FBH1-dependent pathway requires the branch migration activity of RAD54L (3,8,9). Emerging evidence indicates FBH1-mediated fork reversal is mechanistically distinct from the HTLF-dependent pathway. HLTF-mediated fork reversal remodels the fork to form a Holliday-like structure (10,11). In contrast, FBH1 uses its 3’-5’ helicase activity to unwind the lagging strand of a stalled fork to promote a strand annealing independent mechanism (9,12–14).

Once reversed, replication forks are protected from nuclease-mediated degradation by RAD51 and fork protection proteins including BOD1L, FANCD2 and BRCA2 (15–17). Although RAD51 promotes fork protection in both reversal pathways, additional proteins involved in fork protection are specific to fork reversal pathways.

Proteins including BRCA2 and FANCD2 protect replication forks reversed by the HTLF-dependent pathway, whereas BOD1L and FANCA protect forks reversed by FBH1 (3).

Fork reversal restrains replication by potentially error prone DDT pathways that bypass the lesion to continue replication (9,10,18,19). The HLTF-mediated pathway has been shown to restrain replication by limiting PRIMPOL-mediated repriming (10). PRIMPOL is a primase-polymerase that can reprime downstream of a stalled replication fork leaving a ssDNA gap behind the fork. Interestingly, when the HIRAN domain was mutated, unrestrained replication was now mediated by REV1-dependent translesion synthesis (TLS). FBH1 has been shown to restrain replication, but the underlying mechanism has not been determined (9).

In addition to fork reversal, FBH1 cooperates with the endonuclease MUS81 to promote double strand break (DSB) formation and signaling in response to prolonged replication stress (20,21). FBH1 loss is associated with decreased DSB-dependent signaling including pRPA S4/8, pan-nuclear ψH2AX, and pP53 S15 (20–23). FBH1-deficiency results in increased resistance to replication stress including ultra-violet radiation and hydroxyurea (HU) (20–22). The function of FBH1 in the replication stress response has an anti-tumorigenic role protecting melanocytes from UV-mediated transformation (22).

Given increased DDT activity has been associated with increased chemoresistance, we determined how restraining replication by FBH1 contributes to increased resistance to HU. Surprisingly, we found FBH1 restrains replication by both PRIMPOL and TLS in response to HU. However, depletion of PRIMPOL or REV1 did not restore DSB formation in FBH1-deficient cells. We further demonstrate increased activity of PRIMPOL is not preventing replication fork collapse or contributing to replication stress resistance in FBH1-deficient cells. In contrast, REV1 inhibition increased sensitivity of FBH1-deficient cells to hydroxyurea without restoring replication fork collapse. We propose sensitization of FBH1 deficient cells to HU is being driven by REV1-dependent filling of ssDNA gaps left by PRIMPOL. Together, these results indicate FBH1-mediated fork reversal restrains replication by DDT pathways and promotes DSB formation by at least two independent mechanisms.

## Results

### PRIMPOL facilitates unrestrained fork progression in FBH1-deficient cells

Loss of fork reversal in cells deficient of HLTF, SMARCL1, or ZRANB3 leads to unrestrained fork progression during induced replication stress (10,18). FBH1 reverses stalled replication forks and promotes slowing of the replication fork in the presence of HU, but the underlying mechanism was not determined (7). Previous studies have shown PRIMPOL-repriming mediates replication in cells lacking fork reversal mediated by HLTF and SMARCAL1(10,18). To determine whether PRIMPOL was promoting replication in FBH1 KO cells, we used the replication fiber assay to measure DNA tract lengths in the presence of HU in cells after depletion of PRIMPOL (**Fig. 1**). We sequentially pulsed cells with the nucleotide analogs, CldU and IdU. During the secondary IdU pulse we introduced a low dose of HU to slow replication fork progression (**Fig. 1B**). In HU-treated U2OS cells, PRIMPOL knockdown (KD) did not significantly change nascent tract length compared to U2OS cells (**Fig. 1A, 1C).** In FBH1 KO cells, PRIMPOL KD rescued unrestrained fork progression, reducing replication progression to the lengths observed in U2OS cells. We verified this result in RPE-1 cells after co-depletion of PRIMPOL and FBH1 (**Fig. 1D, E**). As observed in U2OS, FBH1-depletion alone resulted in unrestrained replication fork progression in the presence of HU and progression was rescued by depletion of PRIMPOL. Together, these results indicate that FBH1 inhibits unrestrained fork progression by PRIMPOL.

**Figure 1.**
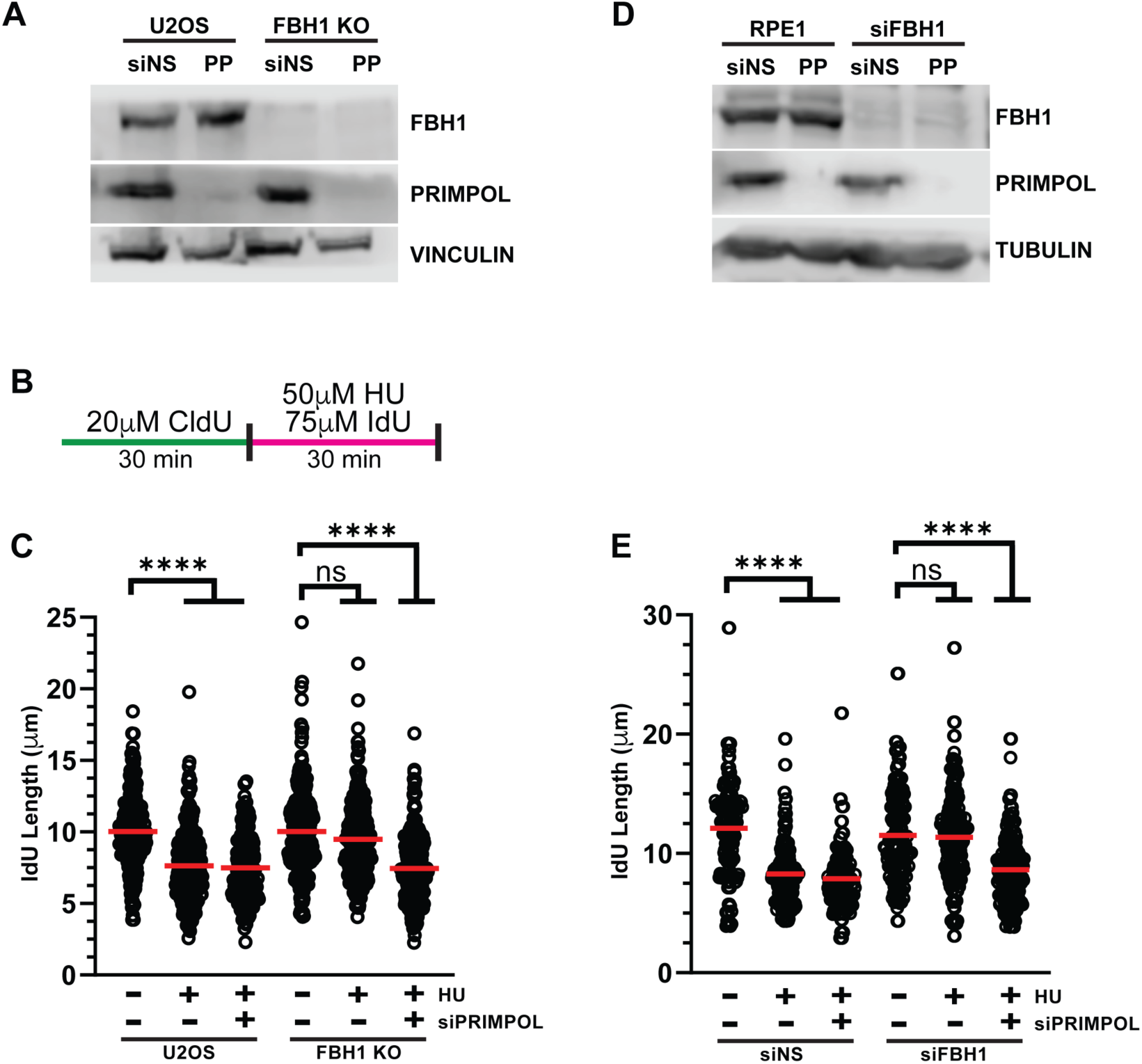
FBH1 inhibits PRIMPOL-mediated DNA synthesis in response to HU. (A) Western blot depicting FBH1 and PRIMPOL protein levels in U2OS cells. Vinculin is the loading control. **(B)** Schematic of the replication fiber experiment. **(C)** Dot plot depicting IdU tract lengths in U2OS cells after the inducted treatments. At least 200 replication tracts per condition. Red line indicates sample mean. **(D)** Western blot depicting FBH1 and PRIMPOL protein levels in RPE-1 cells. Vinculin is the loading control. **(E)** Dot plot depicting IdU tract lengths in RPE-1 cells after the inducted treatments. At least 200 replication tracts per condition. Red line indicates sample mean. n=3, \*\*\*\**p*<0.0001 ANOVA; Tukey HSD

### PRIMPOL does not restore double strand break formation in FBH1-deficient cells

Prolonged replication stress leads to exhaustion of cellular pools of RPA resulting in wide-spread DSB accumulation (i.e., replication fork collapse) known as replication catastrophe (24). Replication catastrophe is marked by high levels of RPA phosphorylation at serines 4 and 8 (pRPA S4/8) and serine 139 on H2AX (ψH2AX). FBH1-deficient cells exhibit reduced DSB accumulation and signaling after prolonged replication stress induced by HU (20–23,25). Previous studies have shown that PRIMPOL-mediated repriming limits ssDNA accumulation and increased PRIMPOL activity is associated with increased chemoresistance in BRCA1-deficient cells (10,18). Thus, we determined whether PRIMPOL-mediated fork progression is resulting in reduced DSB formation and signaling in FBH1 KO cells.

First, we measured the percentage of pRPA S4/8 positive cells after treatment with HU **(Supplementary Fig. 1).** PRIMPOL KD in U2OS cells displayed a slightly higher percentage of pRPA S4/8 positive cells than U2OS transfected with a non-silencing siRNA (siNS). Consistent with previous studies, FBH1 KO significantly reduced the percentage of cells positive for pRPA S4/8. PRIMPOL KD in FBH1 KO cells significantly increased pRPA S4/8 positive cells compared to FBH1 KO alone **(Supplementary Fig. 1B)**. This result suggests PRIMPOL KD is restoring DSB signaling in FBH1-deficient cells.

However, we noticed pRPA S4/8 staining in PRIMPOL KD alone appeared more intense in U2OS or when combined with FBH1 KO. This led us to hypothesize that PRIMPOL was increasing pRPA S4/8 induction independently of FBH1 status. To test this, we measured staining intensity of pRPA S4/8 in RPA positive cells using high-throughput microscopy (**Fig. 2A**). In U2OS cells, PRIMPOL KD resulted in a significant increase in pRPA S4/8 intensity compared to siNS controls indicating PRIMPOL KD is increasing DSB signaling in U2OS cells (**Fig. 2A**). In FBH1 KO cells, we observed a significant decrease in pRPA S4/8 intensity compared to U2OS cells. PRIMPOL KD in FBH1 KO cells slightly increases pRPA S4/8 intensity compared to FBH1 KO alone; however, this increase is consistent with the level observed by knocking down PRIMPOL alone in U2OS cells. We observed a similar result when we co-depleted FBH1 and PRIMPOL in RPE-1 cells (**Supplementary Fig. 2A**). Thus, we conclude that FBH1 KO reduces DSB signaling in PRIMPOL KD and the loss of DSB signaling in FBH1 KO cells is PRIMPOL independent.

**Figure 2.**
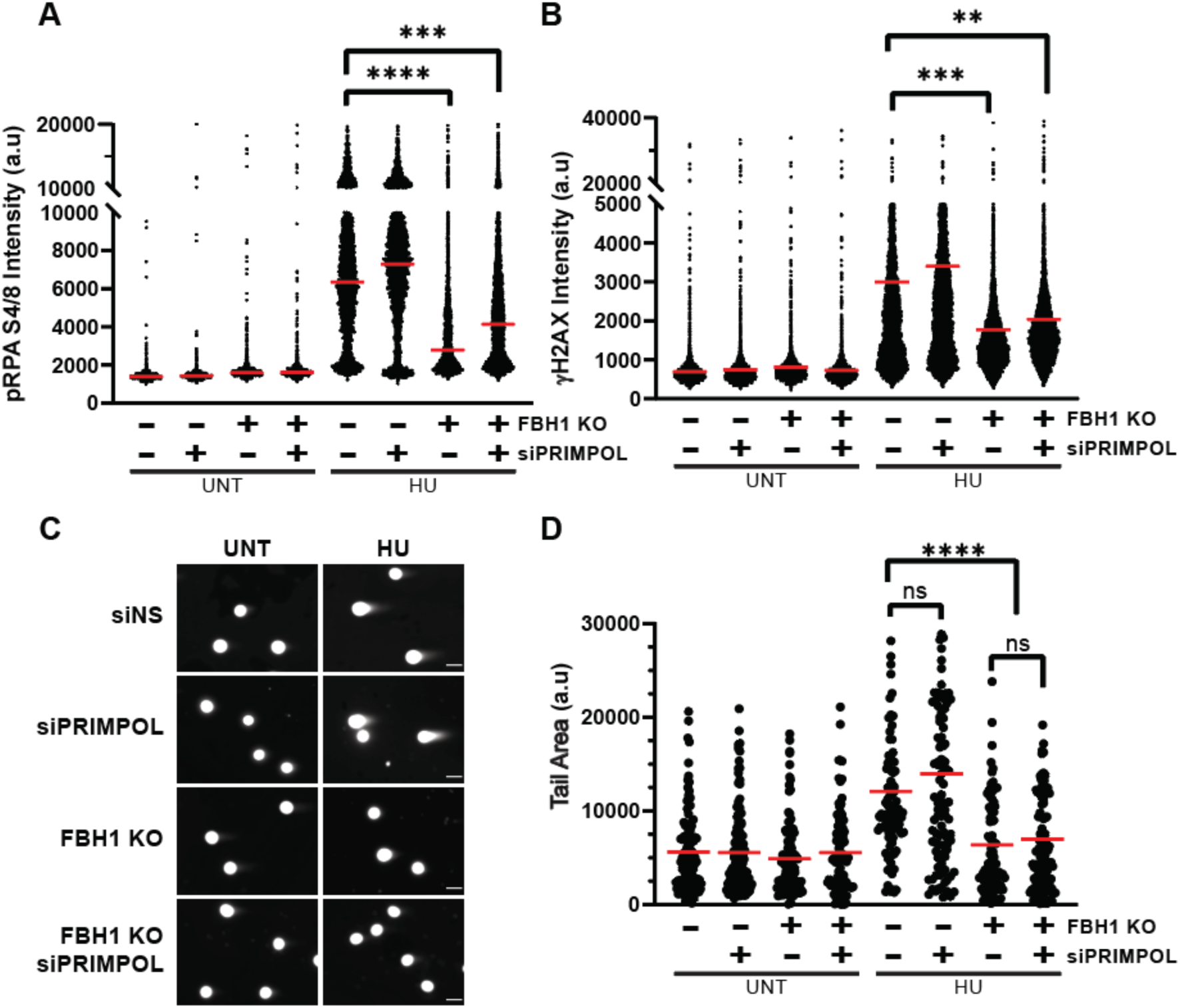
FBH1 KO prevents DSB formation in a PRIMPOL-independent manner. **(A)** Dot plot of at least 3000 nuclei depicting the average pRPA S4/8 intensity in RPA positive cells. Red line indicates sample mean, **(B)** Dot plot of at least 3000 nuclei depicting the average ψH2AX intensity in PCNA positive cells. Red line indicates sample mean **(C)** Representative images of neutral comet assay with DNA stained with SYBR gold (white). Scale bar = 25μm. **(D)** Dot plot of tail area output from neutral comet assay. Red line indicates sample mean. n=3 \*\**p*<0.01 \*\*\**p*<0.001 \*\*\*\**p*<0.0001 ANOVA; Tukey HSD

Next, we looked at ψH2AX intensity in PCNA-positive cells (i.e., S phase) in FBH1 KO cells after PRIMPOL depletion (**Fig. 2B**). As expected, FBH1 KO cells exhibited a significant reduction in ψH2AX intensity compared to U2OS cells alone. Similar to the pRPA S4/8 results, PRIMPOL KD in FBH1 KO cells resulted in a slight increase in ψH2AX, but to a similar extent as observed by PRIMPOL KD alone. We obtained a similar result in RPE-1 cells co-depleted of FBH1 and PRIMPOL **(Supplementary Fig. 2B).** As observed in U2OS, FBH1 KD resulted in a significant reduction in pRPA S4/8 and ψH2AX signaling after HU treatment. PRIMPOL KD resulted in increased pRPA S4/8 and ψH2AX signaling in both U2OS and FBH1 KO suggesting the increase in signaling observed in PRIMPOL KD occurs independently of FBH1 KO. These results indicate the defect in DSB signaling in FBH1-deficient cells is not due to PRIMPOL-mediated DNA synthesis.

FBH1 cooperates with MUS81 to induce DSBs at stalled replication forks after prolonged replication stress (20,21). We determined how PRIMPOL depletion impacted DSB formation using the neutral COMET assay (**Fig. 2C,D**). In U2OS cells, treatment with HU significantly increased DSB formation and PRIMPOL depletion led to a slight, but insignificant increase in DSB accumulation. As expected, FBH1 KO cells exhibited a significant reduction in DSB accumulation compared to U2OS cells (20). In FBH1 KO cells, PRIMPOL KD did not significantly increase DSB accumulation compared to FBH1 KO alone. Together, these results indicated the defect in DSB formation at HU stalled forks in FBH1-deficient cells is PRIMPOL-independent.

### Increased PRIMPOL activity does not contribute to increased cellular resistance of FBH1-deficient cells to HU

FBH1-deficent cells exhibit increased resistance to replication stress induced by HU (20,21). PRIMPOL-mediated DNA synthesis increased resistance of BRCA1-deficient cells to cisplatin (18). To determine if PRIMPOL was leading to increased replication stress resistance in FBH1 KO cells, we measured survival of FBH1 KO cells with and without PRIMPOL KD after increasing concentrations of HU (**Fig. 3**). Compared to U2OS, PRIMPOL depletion slightly increases sensitivity to HU. As expected, FBH1 KO cells exhibited a significantly increase in survival after HU treatment consistent with a role of FBH1 in promoting sensitivity to replication stress. The knockdown of PRIMPOL had no effect on cell survival in FBH1 KO cells after HU treatment. This result indicates that increased PRIMPOL-mediated replication at stalled replication forks is not resulting in increased resistance of FBH1-deficient cells to HU.

**Figure 3.**
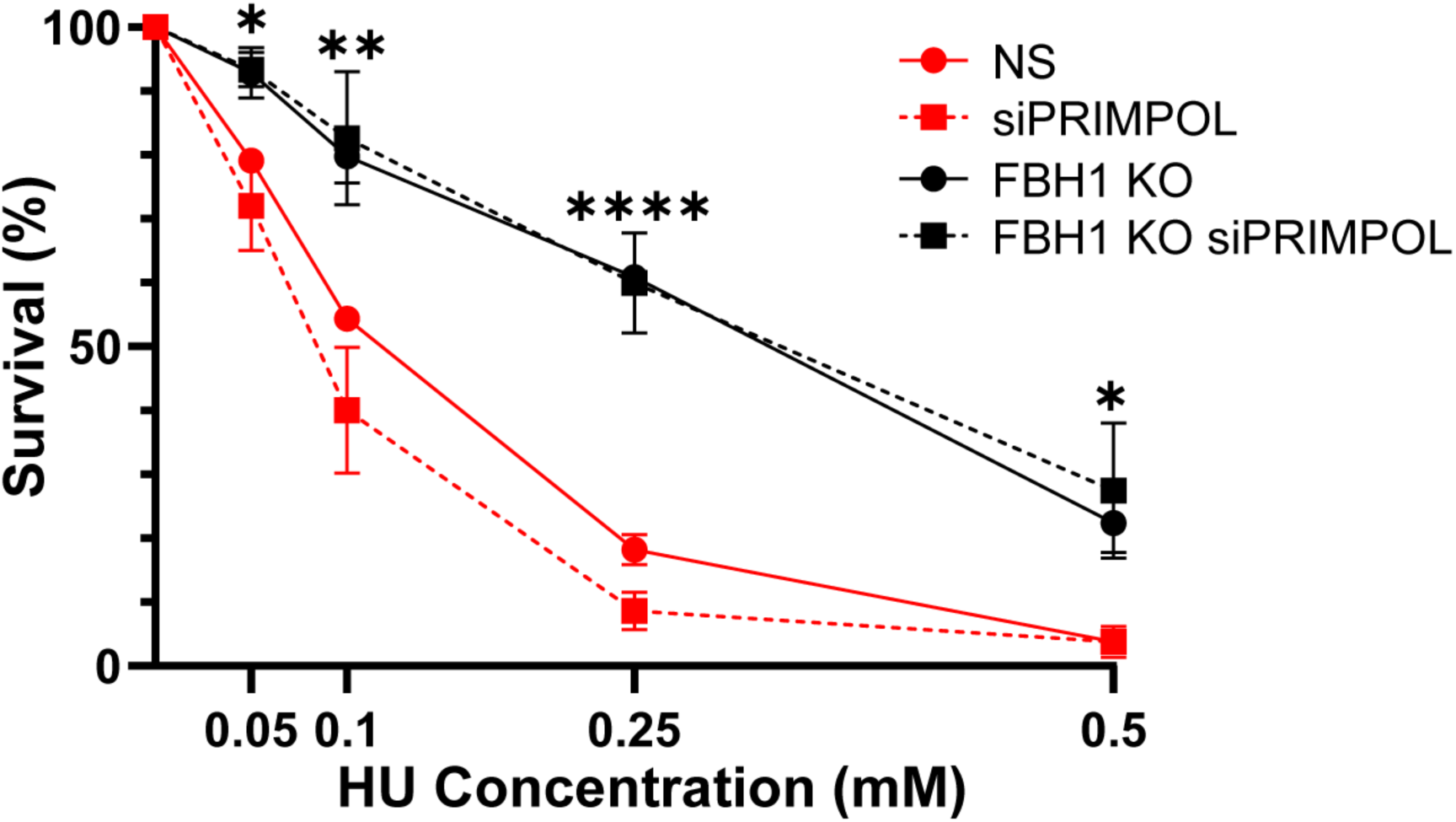
Unrestrained PRIMPOL does not promote increased HU resistance in FBH1 KO cells. Lines represent mean survival after treatment with indicated concentrations of HU. Error bars are SEM. n=3, \**p*<0.05 \*\**p*<0.01 \*\*\*\**p*<0.0001 ANOVA; Tukey HSD

### FBH1 restrains REV1-dependent TLS at stalled replication forks

TLS is a DDT pathway able to bypass DNA lesions and stalled replication forks (26). A previous study discovered that while PRIMPOL was promoting unrestrained fork progression in HLTF KO cells, cells expressing the HIRAN mutant of HLTF limited REV1-mediated TLS at stalled forks (10). Therefore, we hypothesized FBH1 may also inhibit TLS at stalled replication forks. To test this, we examined replication in the presence of HU in U2OS and FBH1 KO cells after treatment with the REV1 inhibitor, JH-RE-06 (hereafter REV1i) (27). In U2OS cells, HU significantly reduced IdU tract lengths compared to untreated cells (**Fig. 4A**). Surprisingly, REV1i rescued unrestrained replication in FBH1 KO cells resulting in reduced IdU tract lengths, similar to HU-treatment in U2OS cells. This result suggests FBH1 also restrains replication mediated by REV1-dependent TLS.

**Figure 4.**
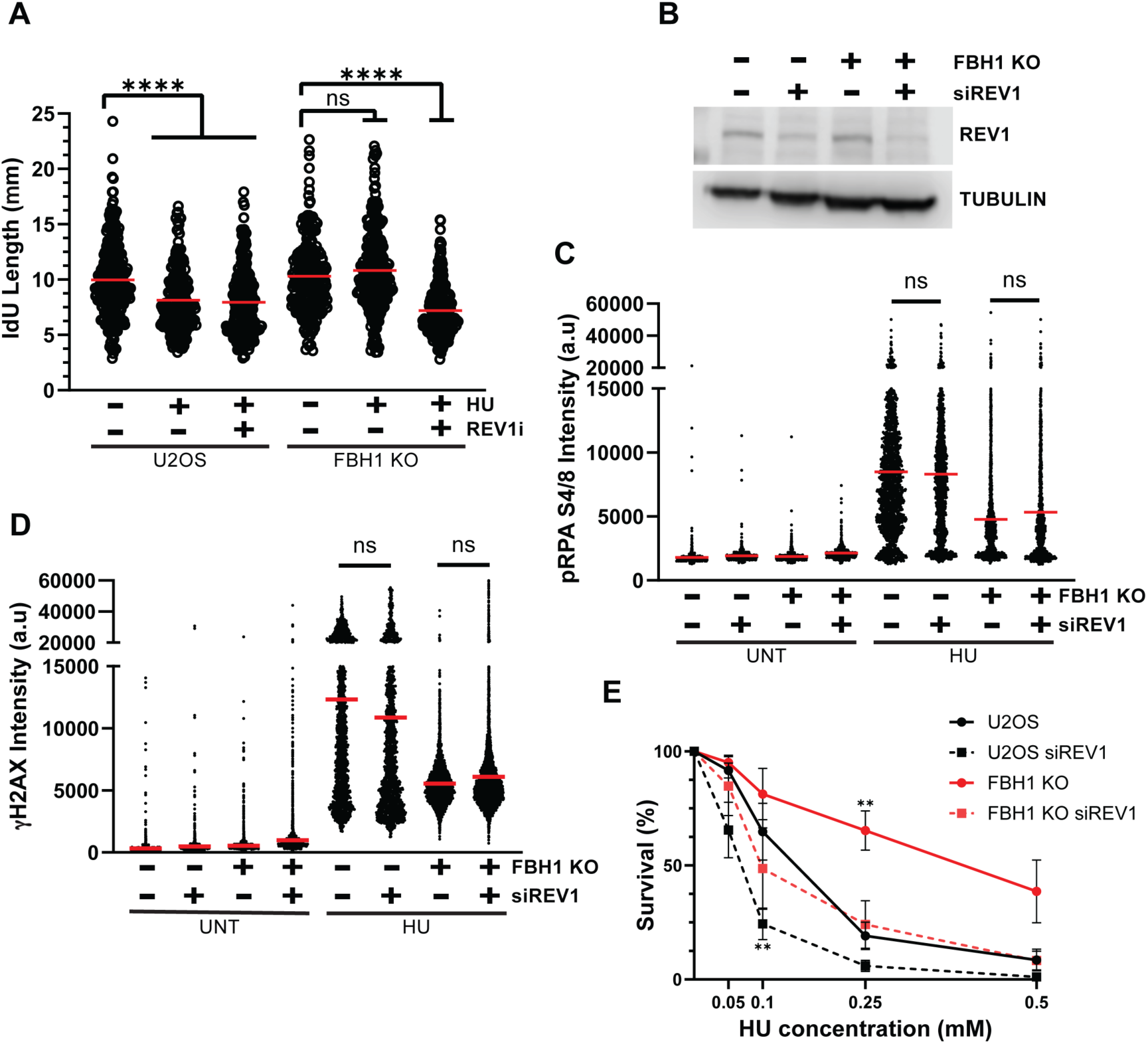
Unrestrained TLS does not promote increased HU resistance in FBH1 KO cells. (**A**) Dot plot depicting IdU tract lengths in U2OS cells after the inducted treatments. At least 200 replication tracts per condition. Red line indicates sample mean. (**B**) Western Blot depicting REV1 protein levels in the indicated samples. Tubulin is used as a loading control (**C**) Dot plot of at least 3000 nuclei depicting the average pRPA S4/8 intensity per cell. Red line indicates sample mean, **(D)** Dot plot of at least 3000 nuclei depicting the average ψH2AX intensity per cell. Red line indicates sample mean (**E**) Line graph depicting percent survival after treatment with indicated concentrations of HU. Error bars are SEM. n=3. \*\**p*<0.01 \*\*\*\**p*<0.0001 ANOVA; Tukey HSD

TLS, like PRIMPOL, is a potentially error-prone DDT pathway and increased activity is known to promote mutagenesis and acquired chemoresistance (28,29). We determined whether REV1-dependent TLS prevented DSB-dependent signaling in FBH1 KO cells by measuring induction of pRPA S4/8 and ψH2AX after treatment with HU. Prolonged exposure to REV1i induced significant DNA damage and cell death, therefore we examined replication stress induction using siRNA transfections to deplete REV1 (**Fig. 4B**). In U2OS cells, depletion of REV1 did not have a significant effect on pRPA S4/8 or ψH2AX induction after HU treatment (**Fig. 4C,D**). Surprisingly, REV1 depletion did not increase pRPA S4/8 or ψH2AX in FBH1 KO cells. These data indicate that REV1-mediated TLS is not responsible for the reduced DNA damage response in FBH1 KO cells.

Finally, we determined whether REV1 contributes to increased HU resistance in FBH1 KO cells by measuring cell survival after treatment to increasing concentrations of HU (**Fig. 4E**). Compared to U2OS, REV1-depleted cells have increased sensitivity to HU. In FBH1 KO cells, REV1-depletion significantly reduced survival after HU treatment, similar to survival observed in U2OS controls. However, REV1-depleted FBH1 KO cells exhibit increased resistance to HU compared to REV1-depletion alone suggesting FBH1 is required to promote hypersensitivity of REV1-depleted cells to HU. Taken together, these data indicate REV1-dependent TLS is not preventing DSB formation and cell death in FBH1-deficient cells.

## Discussion

In this study, we demonstrate that FBH1 restrains replication at stalled replication forks. We demonstrated unrestrained replication is mediated by PRIMPOL and REV1-dependent TLS. In the absence of fork reversal mediated by HTLF, RAD51, and SMARCAL1, replication forks continue to progress due to increased PRIMPOL-dependent synthesis (10,15). Here, we demonstrate that FBH1 limits PRIMPOL-mediated replication at stalled replication forks indicating that restraining PRIMPOL-mediated repriming is a global function of fork reversal. In cells, complete loss of HLTF led to unrestrained replication mediated by PRIMPOL, but when cells expressed a HLTF-HIRAN mutant, unrestrained replication was mediated by REV1-dependent TLS (10).

This result suggests that HLTF localization at the fork, versus loss of fork reversal itself, may be sufficient to block PRIMPOL and promote TLS. In contrast, we demonstrate complete loss of FBH1 results in unrestrained replication by both PRIMPOL and TLS. FBH1 is a 3’-5’ helicase that unwinds the lagging strand to promote fork reversal (12,30). Cells expressing a FBH1-mutant that lacks fork reversal but maintained helicase activity failed to restrain replication (12). Thus, we propose the ability of FBH1 to prevent PRIMPOL-mediated repriming and TLS is likely dependent on formation of the reversed fork. Overall, our results indicate FBH1 regulates the choice between fork reversal, PRIMPOL, and TLS at stalled replication forks.

Unrestrained replication by PRIMPOL and TLS can limit ssDNA accumulation resulting in increased chemoresistance (9,10,18,19). Surprisingly, inhibition of REV1 or PRIMPOL did not restore HU-induced DSB formation and signaling in FBH1-deficient cells. Thus, the ability of FBH1 to restrain replication by TLS and PRIMPOL is independent of the role of FBH1 in promoting MUS81-dependent cleavage.

There are multiple potential mechanisms by which FBH1 can promote replication fork collapse after prolonged replication stress. First, FBH1-mediated fork reversal may be required to generate an intermediate that is susceptible to cleavage by nucleases. Previous work has demonstrated that reversed forks can be processed and cleaved by structure-specific endonucleases (31,32). However, we do not favor this model because loss of the fork reversal proteins, SMARCAL1 and ZRANB3, results in increased sensitivity to a variety of DNA damaging agents including mitomycin C, HU and cisplatin suggesting inhibiting fork reversal alone is not sufficient to prevent replication fork collapse (5,33–35). Second, FBH1 negatively regulates homologous recombination by removing RAD51 from DNA and promoting ubiquitination of RAD51 to prevent rebinding to DNA (23,30,36). Expression of a ubiquitin-resistant RAD51 mutant phenocopied FBH1-deficient cells and exhibited increased resistance to HU (23). However, it is not known how these mutations impact RAD51 biochemical activity, or if other functions of FBH1 contribute to replication stress resistance. Alternatively, FBH1 may prevent replication fork collapse by a mechanism independent of replication fork reversal. FBH1 cooperates with MUS81 to induce DSBs at stalled replication forks (21). FBH1 could be required for efficient localization of MUS81 to the stalled replication fork. Future work will be conducted to determine how FBH1 promotes DSB formation at stalled replication forks.

Despite REV1 and PRIMPOL depletion having no impact on DSB formation and signaling in FBH1-deficient cells, we observed differences in the impact of FBH1 KO to HU sensitivity in PRIMPOL and REV1-depleted cells. HU resistance in PRIMPOL-depleted FBH1 KO cells was indistinguishable from FBH1 KO cells indicating sensitivity of PRIMPOL-depleted cells to HU depends on FBH1. PRIMPOL depletion results in increased ssDNA accumulation in response to HU (18,37). This indicates cell death in PRIMPOL-depleted cells is likely due to FBH1 and MUS81-dependent cleavage of stalled replication forks due to excessive ssDNA accumulation. We demonstrate REV1-depleted cells are hypersensitive to HU and FBH1 KO increases resistance to levels observed in U2OS siNS controls. Thus, there are two mechanisms contributing to cell death in REV1-depleted cells: FBH1-dependent replication fork collapse and a mechanism that is FBH1-independent (38).

REV1-dependent TLS occurs at multiple locations in response to replication stress. At a stalled replication fork, “on-the-fly” TLS on the leading strand bypasses lesions preventing replication fork collapse (39,40). We propose FBH1 KO increasing resistance of REV1-depleted cells to HU is likely to loss of DSB formation at stalled forks that arise due to inhibition of TLS at the stalled replication fork. TLS also fills ssDNA gaps post replication and during mitotic DNA synthesis (MiDAS) to prevent cells with under-replicated DNA from undergoing mitotic division (38,41,42). Unrestrained replication by PRIMPOL results in ssDNA gap accumulation that are filled, in part by TLS, prior to mitosis. We propose ssDNA gap accumulation due to PRIMPOL-mediated synthesis is killing FBH1 KO cells due to cells undergoing mitotic division with under-replicated DNA. Two lines of evidence support this model. First, FBH1 KO cells exhibit increased 53BP1 bodies in G1 phase of the cell cycle after recovery with HU indicative of regions of under-replicated DNA (7). Second, FBH1 KO cells are sensitized to WEE1 inhibitors despite preventing replication fork collapse due to increased mitotic catastrophe (25).

FBH1 loss has been linked to melanoma and mutation or loss of FBH1 may be present in additional tumors (22,43). An increase in PRIMPOL-mediated synthesis or TLS would give FBH1-deficient cells a proliferative advantage by allowing cells to continue replication in the presence of DNA damage. However, DDT up-regulation may be a mechanistic vulnerability in FBH1-deficient tumors. Specifically, targeting DNA gaps left by PRIMPOL may be a viable chemotherapeutic strategy to kill FBH1-deficient tumors. Single-stranded gap accumulation by PRIMPOL requires gap filling by template switching or TLS in late S-phase and G2 before mitotic entry (42). The accumulation of gaps in BRCA-deficient cells contributes to cisplatin sensitivity (44). Furthermore, TLS reduces replication stress in cancer cells by gap suppression and tumors become dependent on TLS for survival (45). In TLS-dependent cancer cells, TLS inhibitors synergized with inhibitors that induce replication fork collapse such as ATR or WEE1 inhibitors. Given FBH1 KO cells are sensitized to WEE1 inhibition, an interesting future area of study would be to determine whether combinatorial inhibition of WEE1 and TLS is lethal to FBH1-deificient tumors (25).

In conclusion, we demonstrate FBH1 restrains replication by PRIMPOL and TLS in response to HU-induced replication stress. However, inhibition of PRIMPOL or TLS does not restore DSB formation in response to HU suggesting FBH1 restrains replication and promotes replication fork collapse by two independent mechanisms. Finally, we demonstrate inhibition of REV1, but not PRIMPOL, increases sensitivity of FBH1 KO cells to HU suggesting FBH1 KO cells may be dependent on gap filling mechanisms to tolerate replication stress. Additional work being conducted will determine whether targeting TLS inhibition alone or in combination with WEE1 inhibition can be exploited as a chemotherapeutic strategy to kill FBH1-deficient tumors.

## Materials & Methods

### Cell culture

U2OS cells were grown in DMEM (Gibco, 1196518) supplemented with 10% FBS (Atlas Biologicals, S10350H). hTERT immortalized RPE1 cells were grown in DMEM/F12 (Corning, 10-092-CVR) supplemented with 10% FBS. Cells were grown at 37°C in 5% CO2. HU and cisplatin were resuspended in water and diluted in media at the indicated concentrations.

### siRNA transfections

Cells (1x10^5^) were seeded in 6-well plates and allowed to adhere overnight. Transfections were prepared in 1x Opti-MEM (Gibco) and Lipofectamine RNAiMAX (Fisher Scientific, 13778150) following manufacturer instructions. Two siRNAs against FBH1 (Thermo) were pooled and transfected at a final concentration of 75μM. The siRNA targeting PRIMPOL and REV1 was from Dharmacon (PRIMPOL: J-015804, REV1 (smartpool consisting of siREV1 1-4): L-008234-00). The All Star Negative control siRNA (Qiagen) was used a non-silencing (siNS) control. siRNA sequences are:

siFBH1-1 5’ GGAGTCAGGAGACCCGGGTTCTTGT

siFBH1-2 5’ CCGGGTTCTTGTGTTGGCACCATCAC

siPRIMPOL 5’ GAGGAAAGCUGGACAUCGA

siREV1-1 5’ GGACAAGGACCGUAAACGA

siREV1-2 5’ CAGCUUAUCUCCUGUCGA

siREV1-3 5’ GCACAAAGAACGAGAACGA

siREV1-4 5’ CCAACUUUGAGUCGGGAAA

### Immunofluorescence

Cells (4x10^4^) were seeded in 12-well plates with 22mm circular cover slips and allowed to adhere overnight. Cells were treated with 2mM HU for 24 hours. Cells were permeabilized a HEPES/TritonX-100 buffer (20mM HEPES, ph7.4, 3mM MgCl2, 50mM NaCl, 0.5% Triton X-100) for ten minutes and fixed to the coverslips with ice-cold methanol. Coverslips were incubated in primary antibodies overnight at 4°C. Primary antibodies used were PCNA (ABCAM, ab18197, 1:1000), ψH2AX (ABCAM, ab26350, 1:1000), RPA (ABCAM, ab2175, 1:1000), pRPA S4/8 (Bethyl Laboratories, A300-245A- M, 1:1000). Cells were washed three times in 1X PBS before staining with secondary antibodies. Coverslips were incubated in Alexa fluor-conjugated secondary antibodies (Invitrogen, 1:1000) for one hour at room temperature. Coverslips were washed 3 times for five minutes in 1X PBS followed by two minutes each in 70%, 95% and 100% ethanol and allowed to air dry. Cells were mounted on slides with vectashield containing DAPI. Cells were imaged on ImageExpress Micro 4 (Molecular Devises) and analyzed using MetaXpress software version 6.7.2.290.

### Replication fiber assay

Cells (4x10^4^) were seeded in 12-well plates and allowed to adhere overnight. Cells were pulsed with 20uM CldU for 30 minutes. Cells were quickly washed with 1X PBS and incubated in 75uM IdU with or without 50uM HU for 30 minutes. For assays with REV1i cells were incubated with REV1i , JH-RE-06, for 30 minutes prior to the first pulse and REV1i remains throughout the subsequent pulses. Cells were collected by trypsinization and resuspended in ice-cold 1X PBS at 4X10^4^ cells/ml. Replication fibers were prepared as previously described (46). Slides were stained with mouse BrdU (BD, 347580, 1:50) to label IdU and rat BrdU (ABCAM, ab6326, 1:200) to label CldU. Images were acquired at 60x magnification using a Zeiss Imager.M2 epifluorescence microscope with an Axiocam 503 mono camera. The length of at least 150 replication tracts was measured using ImageJ.

### Neutral comet assay

Cells (2x10^5^) were seeded in 6 well plates and allowed to adhere overnight. Media was refreshed with or without 2 mM HU for 24 hours. Cells were collected and suspended in 500 μl PBS. Neutral comet assay was conducted as described in the following paper excluding lysis buffer and comet slides that were purchased from R&D Systems (4250-050-K) (47). Comets were acquired at 20x magnification using a Zeiss Imager.M2 epifluorescence microscope with an Axiocam 503 mono camera. Tail area was calculated using the open comet plugin in ImageJ (48).

### Survival assay

Cells (1x10^3^) were plated in triplicate in black walled 96 well plates (Corning, 3606) and allowed to adhere overnight. Cells were treated with HU for 48 hours and allowed to outgrow in fresh media for 7 days. After outgrowing live cells were stained with Hoechst 33342 (1:2000) for 20 minutes and counted using the ImageExpress Micro 4.

### Generation of FBH1 polyclonal antibodies

The N-terminal 250 amino acids of human FBH1 were cloned in to a pGEX6p-1 expression vector using Gibson assembly. Primers used to amplify the FBH1 fragment were FBH1_pGEX6_F (5’ TTCTGTTCCAGGGGCCCCTGGGATCCagctacgaggtgacttcag) and FBH1_N250_pGEX R (5’ GTCGACCCGGGAATTCCGGGGATCCtcaaagcccatagtatgagtcag). GST-FBH1[1-250] in the pGEX6P-1 vector (Amersham/GE Healthcare/Cytiva) was introduced into BL21:DE3 Rosetta cells (Novagen). The cells were grown at 37 °C to an OD600 of 0.8 and induced with 0.4 mM IPTG for 16 h at 16 °C. All the steps of the purification were performed at 4 °C. A 10 g pellet of cells was suspended in 50 mL of cell breakage buffer (50 mM Tris-HCl, pH 7.5, 10% sucrose, 2 mM EDTA, 250 mM KCl, 0.01% IgePal, 1 mM DTT, 1 mM Phenylmethulsulfonyl fluoride, 0.5 mM benzamidine and 5 μg/mL each of aprotinin, chymostatin, leupeptin, and pepstatin). The cells were sonicated, and the crude lysate was centrifuged at 100,000 x g for 90 min. The clarified lysate was incubated with 1 mL of glutathione Sepharose (Cytiva) with constant agitation. The matrix was poured into a 1 cm column and washed with 10 mLs Buffer A (50 mM Tris-HCl, pH8.0, 10% glycerol, 1 mM EDTA, 1 mM DTT, and 0.01% IgePal) containing 1 M KCl and then 5 mL of Buffer A containing 300 mM KCl. The bound GST-FBH1[1-250] was eluted with 6 mL of 10 mM glutathione in Buffer A containing 300 mM KCl. The fractions containing GST-FBH1[1-250] were pooled and concentrated in an Amicon Ultra microconcentrator to 500 μL and subjected to size exclusion chromatography on a 40 mL Sephacryl S200 column in Buffer A containing 300 mM KCl. The fractions containing GST-FBH1[1-250] were pooled and concentrated as before to 8 mg/mL, and stored in small aliquots at -80 °C. ∼1 mg of GST-FBH1[1-250] was used to generate antibodies from Pacific Immunology. Purified GST-FBH1[1-250] was coupled to CNBr activated Sepharose per the manufacturer’s instructions. Serum was applied to a 1 mL of the matrix in a 1 cm column. The matrix was washed with 10 mL PBS (10 mM Na2HPO4, pH 7.4, 2.7 mM KCl, 137 mM NaCl, 0.05 % TWEEN) followed by elution with 166 mM MES pH 6.0 containing 100 mM diethylamine. The fractions that contained the antibodies were pooled and dialyzed against PBS. The dialyzed antibodies were concentrated in an Amicon Ultra microconcentrator, supplemented with 0.01% sodium azide and stored in small aliquots at 4 °C.

### Western Blotting

Whole cell extracts were prepared by resuspending cells (10^6^/100 μl) in Laemmli buffer (62.5mM Tris-HCl, ph6.8, 2% SDS, 10% glycerol, 5% 2-mercaptoethanol, and 0.002% bromophenol blue). Protein extracts were run on an SDS-Page gel and transferred to a PVDF membrane. The membranes were blocked and incubated with antibodies diluted in 5% milk. REV1 westerns were blocked and incubated in 5% BSA instead of milk. The membrane was stained with primary antibodies recognizing FBH1(1:1000 Pacific immunology: purification described above), PRIMPOL (kind Gift from Juan Mendéz (37)), REV1 (Santa Cruz, sc-393022, 1:100), and Vinculin (Cell signaling, 13901S, 1:1000) overnight at 4°C. The membrane was washed 3X with TBS + 0.1% Tween-20 and then incubated with HRP secondary antibodies for 1 hour at room temperature (LI-COR, 926-80011/10, 1:2000). Membranes were incubated with WesternSure Premium Chemiluminescent substrate (Li-COR) following manufacturers’ instructions. Membranes were imaged and analyzed using the LI-COR C-DiGit imager and Image Studio software version 5.2.5.

### Statistical Analysis

All experiments were conducted at least three independent times unless otherwise indicated. Statistical analyses were performed using Graphpad Prism (version 10.2.3). Significance was calculated using an ANOVA followed by Tukey HSD. Significance for the pRPA S4/8 and ψH2AX intensity data was conducted on the means of each experiment. Statistical significance was determined by a *p*-value below 0.05.

## Author Contributions

JLT and JMM designed the research study. JLT, GM, SLR, TJM, and ZRT conducted the experiments and analyzed data. JLT and JMM wrote and edited the manuscript.

## Data availability

Further information and requests for resources and reagents should be directed to and will be fulfilled by the lead contact, Jennifer Mason, PhD.

## Acknowledgements

We would like to thank Garrett Buzzard, Amarachukwu Onoh, and Michael Sehorn for purification of GST-tagged FBH1-N250 and FBH1 polyclonal antibodies.

