## Supplemental Figures for "FBH1 restrains replication progression independently of promoting double strand breaks at hydroxyurea-stalled replication forks"

### Supplementary Figures

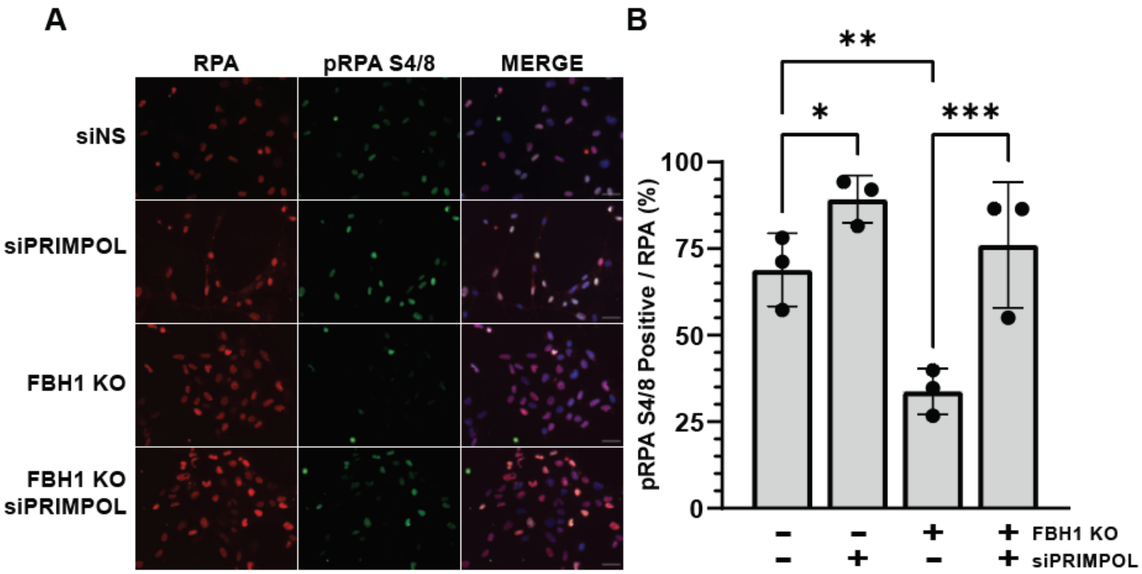


**Supplementary Figure 1. PRIMPOL depletion increases percentage of pRPA S4/8 positive cells in FBH1 KO cells. (A)** Representative images of total RPA (RED), pRPA S4/8 (GREEN), and merged with DAPI**.** Scale bars are XXX µm **(B)** Bar graph of low depicts percent of pRPA S4/8 positive cells out of total RPA positive cells. Error bars are standard deviation. *$p$<0.05 **$p$<0.01, ***$p$<0.001 ANOVA; Tukey HSD

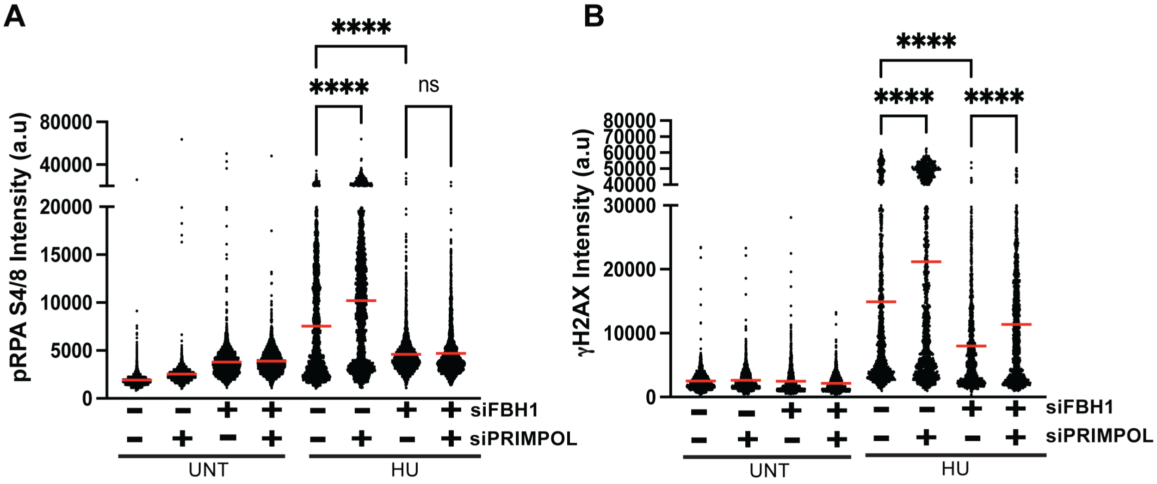


**Supplementary Figure 2. PRIMPOL depletion does not restore DSB signaling in FBH1-depleted RPE-1 cells after HU treatment. (A)** Dot plot depicting average pRPA S4/8 intensity of at least 3000 cells. Red line indicates sample mean. **(B)** Dot plot depicting average γH2AX of at least 3000 cells. Red line indicates sample mean. n=2. Both independent experiments had the same result. ns, not significant; ****$p$<0.0001 ANOVA; Tukey HSD
